# On whole-genome demography of world’s ethnic groups and individual genomic identity

**DOI:** 10.1101/2022.03.28.486119

**Authors:** Byung-Ju Kim, JaeJin Choi, Sung-Hou Kim

**Affiliations:** Department of Chemistry and Center for Computational Biology, University of California, Berkeley, CA, 94720, USA; Human Genome Research Center, Incheon National University, Incheon 22012, Republic of Korea (South Korea); Division of Biological Systems and Engineering, Lawrence Berkeley National Laboratory, Berkeley, CA, 94720, USA; Department of Molecular Biophysics and Integrated Bioimaging Division, Lawrence Berkeley National Laboratory, Berkeley, CA, 94720, USA; Convergence Research Center for Insect Vectors, Incheon National University, Incheon 22012, Republic of Korea (South Korea)

**Author notes:** Corresponding author: Sung-Hou Kim.

## Abstract

All current categorizations of human population, such as ethnicity, ancestry and race, are based on various selections and combinations of *subjectively-* and/or *qualitatively*-defined characteristics, such as ancestral lineage/location, cultural/societal norm, language, skin color and other phenotypes and traits perceived by the members within or from outside of the categorized group. Yet, such categorization has been broadly used also in the fields of human genetics, health sciences and medical practices (e.g., *1,2,3*), where the observed health characteristics are objectively and quantitatively definable, but the population categorization is not yet available. Here we show the feasibility of deriving a whole-genome-based categorization that is objectively definable and quantitatively measurable. We observe that: (a) the world’s ethnic populations form about 14 genomic groups (GGs); (b) each GG consists of multiple ethnic groups (EGs); and (c) at an individual level, approximately 99.8%, on average, of the whole genome contents are identical between any *two individuals* regardless of their GGs or EGs.

## Introduction

### Background

Historically, classification or categorization of human population, such as race, ethnicity, and ancestry has been attempted by using various physical, cultural and sociological characteristics, subjectively-presumed ancestry, language, cultural history, religions, socioeconomic status and others. Such categorizations and characteristics are, in general, difficult to define objectively and quantitatively. Therefore, in this era of genomics, they are, in general, unsuitable for correlating between the categories and the characteristics of whole-genome data gathered objectively and quantitatively in the fields of human genetics, health sciences and medical practices. Thus, there is an urgent need for a objectively and quantitatively definable categorization of extant human population that can be used to correlate with genomic characteristics in these fields (*1, 2, 3*).

Several large-scale genomic studies have been published during the last decade to address the extent and types of genomic diversity of the extant human species, e.g., for 2,504 individuals from 26 large “population groups (PGs)” from diverse geographic locations in the 1000 Genomes Project (1KGP) (*4*), for 300 individuals from 142 ethnic groups (EGs) based on ancestry, linguistic, faith and cultural differences in the Simons Genome Diversity Project (SGDP) (*5*), and for 44 African ethnic populations in 4 linguistic groups (*6*). As a result of these studies as well as other similar or related studies (e.g., *7, 8*), a large body of whole-genome Single Nucleotide Polymorphism (SNP) data became publicly available for a wide range of individuals representing different ethnic groups or “population” groups throughout the world.

Therefore, we revisit these data for the purpose of (a) finding a whole-genome-based categorization for the ethnic groups in these studies using a text-comparison method of Information Theory (*9*), which has not been applied in any of the earlier studies, and (b) quantifying the fraction of whole genome that enables such genome-based categorization.

### Objectives

The itemized objectives of this study are two folds: First, (a) using the concept of the “contextually-linked Single Nucleotide Variation (c-SNV) genotypes”, identify whole-genome-based genomic groups (GGs) of the world’s EGs in the SGDP database (*5*), where c-SNVs are defined by overlapping short strings of ordered SNV genotypes, (b) identify a set of EGs that belong to each GG, (c) suggest the order of emergence of GGs; and Second, (d) quantify the magnitudes of genomic identity/difference between two individuals within each GG as well as between two different GGs to estimate the fraction of whole genome that contributes toward an individual’s GG; and (e) perform a similar study as (d) above for the 1KGP samples, which has fewer EGs divided into 26 broadly defined “Population Groups(PGs)”, but with larger sample size.

### Approach

Our approach is to adapt a computational method of comparing and categorizing linear information such as texts or books in the field of Information Theory (*9*) by “word frequency profiles” (*10*). We generalized the method to be applicable to other linear information, such as genomic sequence, proteomic sequence, or SNV sequence. In this study we convert the whole-genome SNV genotypes of an individual into a “book” of “Feature Frequency Profile (FFP)” (*11*), where each “Feature” (which corresponds to a unique “word” in a book) consist of a *unique* c-SNV as a “character” of the genomic feature, and its frequency in a genome, as the Feature’s “character state”, for a given length of c-SNVs (see ***Feature Frequency Profile*** in Data source and Methods of Supplementary Materials). Thus, an FFP of an individual’s c-SNV genotypes and their frequencies contains all the information necessary to reconstruct the original sequence of the ordered SNV genotypes. Then, for a given length of the c-SNVs, all pair-wise divergence of the FFPs of study population can be calculated to assemble a “genomic divergence matrix” for the given c-SNV length. This matrix is used both for Principal Component Analysis (PCA) (*12*) to discover the genome-based demographic grouping pattern (as manifested by clustering) of the study population (see **FFP-based *PCA*** in Data Source and Methods of Supplementary Materials) and to find the genome-based clading pattern from a rooted neighbor-joining tree (*13*; see ***FFP-based rooted NJ tree*** and ***Tree rooting*** in Data Source and Methods of Supplementary Materials). Among many such trees, each corresponding to a given length of the c-SNVs, one with the most topologically stable tree is selected as the final tree. From the final tree the order of the emergence of the founder nodes of GGs are also predicted on an evolutionary progression scale, which corresponds to the normalized cumulative branch-lengths from the tree root to each of the founder nodes of the GGs.

The uniqueness of our approach is two folds: we compare the entirety of the individual’s whole genomic variations *in context* and each individual’s whole genome variation is compared to that of each of all other individuals, not to that of one Eurocentric “human reference genome”.

## Results

Our results are divided into two parts: the first part focuses on, at the group level, the whole-genome-based grouping pattern of the population of 164 ethnicities from the recent SGDP database, and on the emergence order of the founders of the groups. These provide the results for the first three specific objectives ((a) – (c)) listed under **Objectives** in the Introduction. The second part focuses, at the individual level, on the extent of genomic identity at all SNP loci between two individuals among all members of two databases: the SGDP sample, which has the largest number of ethnic groups, and the 1KGP populations, which has the largest number of samples per population group. Then, we extrapolate the percent identity of all SNP loci to that for the respective whole-genome length to get an estimate of intuitively understandable magnitude of the whole-genomic identity between two human individuals within a given GG or PG and between two different GGs or PGs. These provide the results for the remaining two specific objectives ((d) and (e)) listed under **Objectives** in the Introduction.

### Part I. Genomic demography: 14 “genomic groups (GGs)” and the order of their emergence

At a group level, we examined the genome-based grouping patterns and their relationships among all GGs by two different methods, both based on the individual’s genomic variation expressed by the FFPs of c-SNVs: Principal Component Analysis (PCA) (*12*) and Neighbor-Joining (NJ) phylogeny methods (*13*).

### Clustering by PCA

Figure 1 shows the grouping pattern, based on clustering revealed by PCA for all members (345 individuals) of the study population from 164 EGs in the recent SGDP database (*5*), where the genomes were sequenced to an average coverage of 43-fold. In the PCA, the clustering is strictly based on the genomic divergence matrix calculated using FFPs of c-SNVs (see ***FFP based PCA*** in Data source and Methods of Supplementary Materials). The figure shows that:

**Figure 1.**
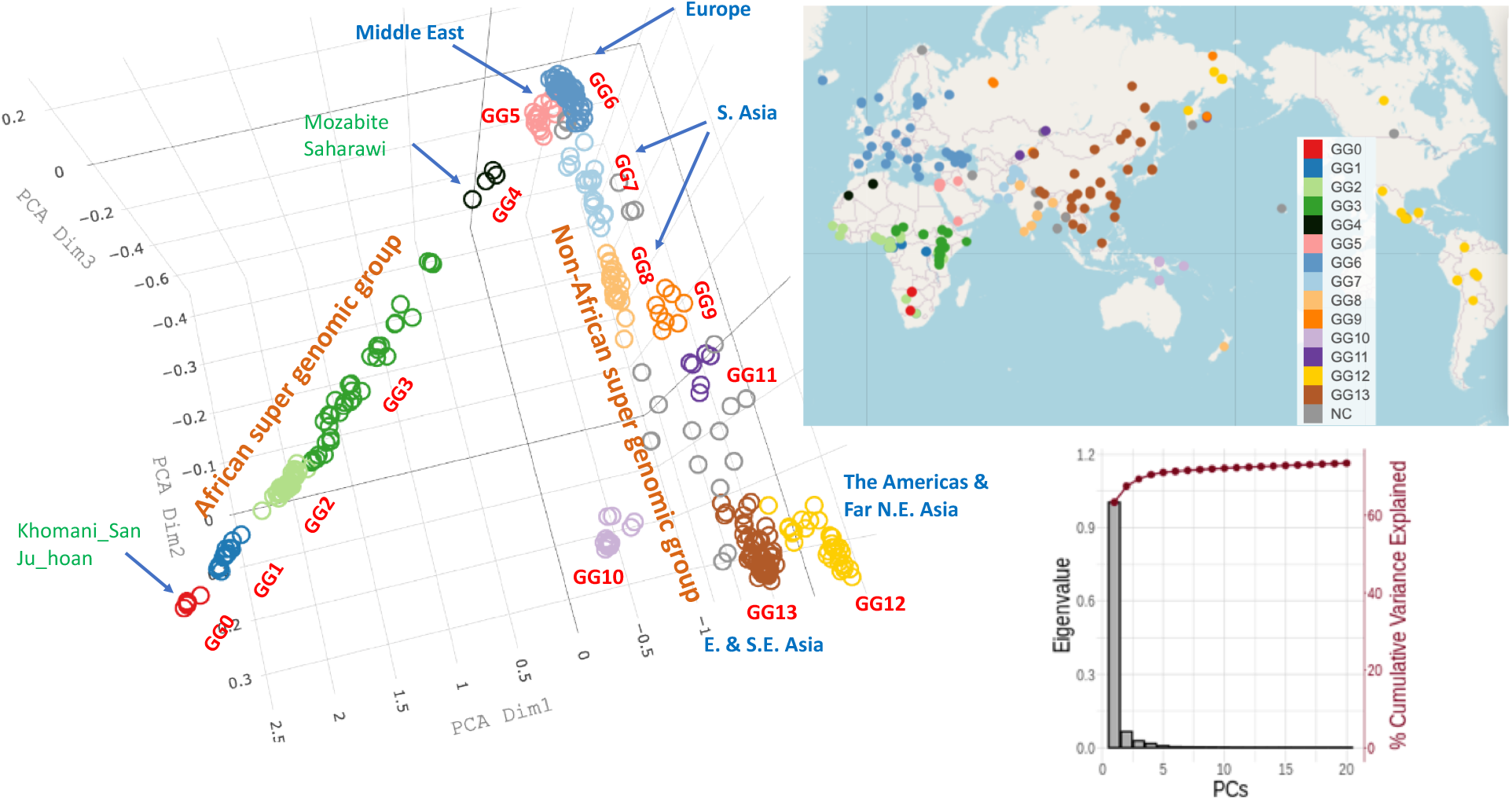
Clustering pattern by genomic-divergence-based PCA plotted for the three major principal component axes. PCA was performed using the “distance matrix”, where each “distance” is calculated by JS divergence between two FFPs of c-SNVs. Most of 345 individuals from 164 ethnic groups from the updated SGDP form about 14 clusters shown in different colors and labeled as genomic groups (GGs) GG0 – GG13, of which most are tight clusters, but some are not, e.g., GG3 consists of multiple un-resolved clusters of African ethnic groups. All un-clustered individuals are shown as gray circles and located outside of Africa. Although the EG members of each GG are tightly clustered in genomic divergence space (PCA space), they are widely spread out within a large region bound by great geological barriers, as shown at the geographical locations of the sample collection sites, suggesting broad migration of the EGs within the region (see the top right inset). Sorted eigenvalues and variance explained in % for the first 20 principal component axes are shown in the bottom right inset. Grouping of GGs is based on a combination of the clustering pattern of individuals in PCA (in genomic divergence space), the clading pattern in NJ tree (in genomic divergence space), and the clustering pattern in geological/geographical map. A video of the rotating PCA plot is shown in Supplementary Fig. S1A.

a. There are about 14 clusters in genomic variation space (i.e., c-SNV space) which we named “Genome-based Groups” or “Genomic Groups (GGs)” (one of which, GG3, consists of a collection of several not-well-resolved sub-clusters of various sizes);
b. All GGs are divided into two interconnected super groups: one defined by a long linear arm in Fig. 1 containing all five African GGs (GG0 – GG4), most of which are linearly linked but not well clustered (due to sparse availability of the whole-genome sequences representing the vast diversity of African EGs) and account for all 45 African EGs available in the SGDP database. The other is a more fanned-out arm containing all non-African GGs (GG5 – GG13), most of which are well clustered, accounting for all of the 119 non-African EGs;
c. Most of African GGs are not well resolved and linearly connected, with GG0 (consisting of the EGs of Khomani_San and Jo_hoan) at one end, and GG4 (consisting of the EGs of Mozabite and Saharawi) at the other end, but all non-African GGs appear to have originated from the Middle East GG (GG5). Most of non-African GGs are tightly-clustered and well-resolved, but, all remaining un-clustered and isolated samples are found in this super group;
d. Each GG consists of multiple EGs (see Supplementary Table S1), thus there is no one-to-one correspondence between GGs and EGs;
e. Many of the 14 GGs can be assigned to various geographical or geological regions (see the color coding in the upper inset of Fig. 1);
f. The extant members of some large GGs (GG6, GG12 and GG13) are widely scattered in geographical space (see the upper inset of Fig. 1), but their genomes are closely clustered in genomic variation space (see the PCA plot of Fig. 1 and Fig. S1A in Supplementary Materials).

Some of the features described above (such as (b), (c) and (f) mentioned above) have been observed in the first SDGP publication (see Extended Data Fig. 4A of reference *5*). We also noticed a few unexpected observations in our PCA plot where some members of a given ethnic group are found in two geographic locations very far apart (see **Supplementary Note 1** in Supplementary Materials).

### Clading in neighbor-joining (NJ) tree

Figure 2 shows the clading pattern revealed in our rooted FFP-based tree with cumulative branch-lengths (see ***FFP-based rooted NJ tree*** in Data source and Methods of Supplementary Materials) by a neighbor-joining method (*13, 14*) (for a corresponding topological tree with the names of EGs and clading details is shown in Supplementary Fig. S1B, which can be expanded for easier viewing). The genomic divergence matrix used as the input of the tree is the same as that used in the PCA clustering for Fig. 1 above, but, in the NJ tree-building method, two additional conditions are imposed for the evolutionary model, which constrains the tree topology and clading: (a) bifurcating branches emerging from each internal node and (b) maximum parsimony (minimal evolution) when choosing the neighbors to join as a sister pair. For comparison, all the GGs identified by the PCA clustering in Fig. 1 are also shown in the middle circular band of Fig. 2. Noticeable from the tree are:

**Figure 2.**
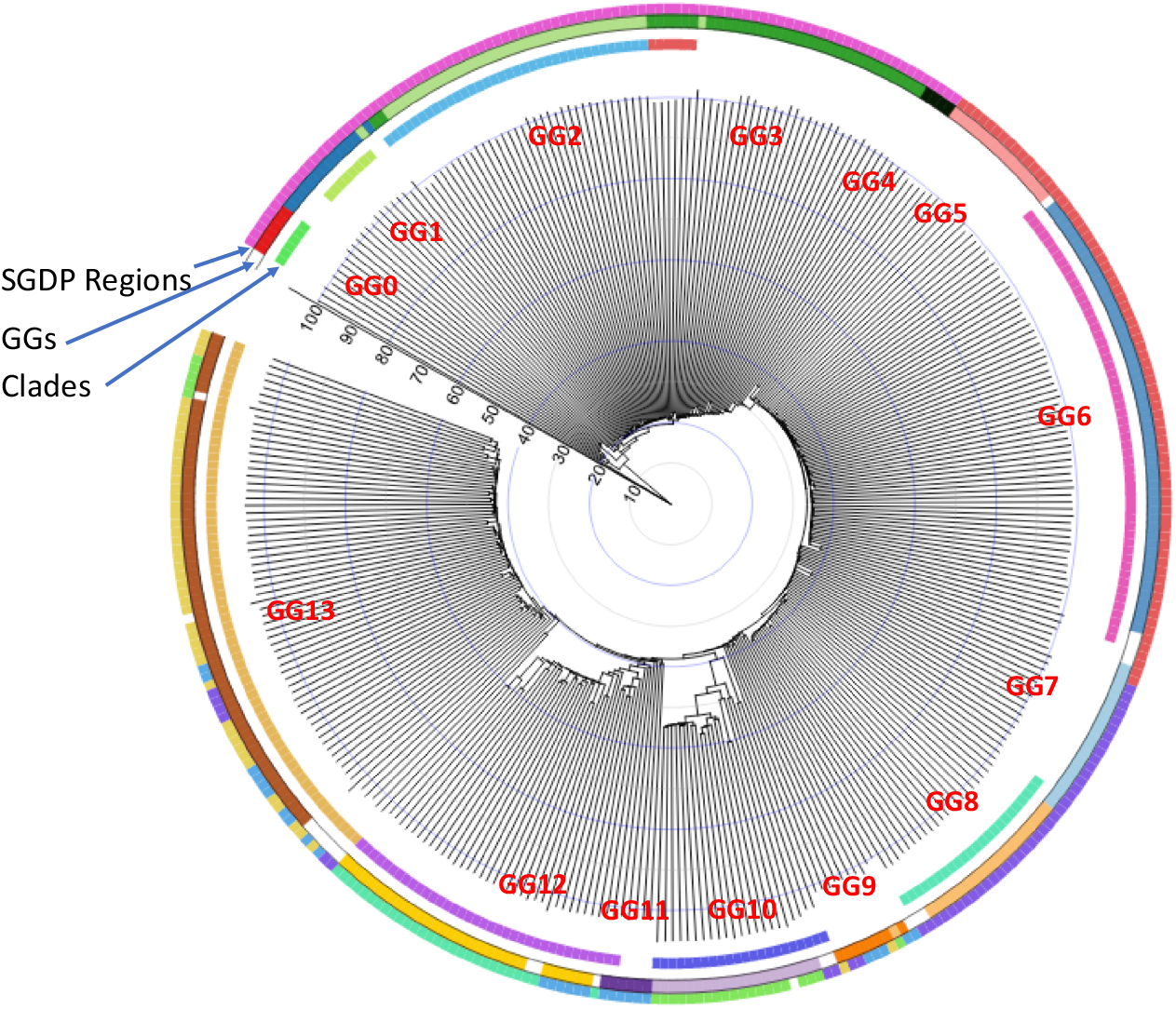
Rooted c-SNV-based NJ tree in circular form (*21*) with the “evolutionary progression scale (EPS)”. The same “distance matrix” as was used in Fig. 1 is used to construct the tree (see **Rooting of c-SNV-based tree** in Data Source and Methods in Supplementary Materials). “Evolutionary progression scale” is the scaled cumulative branch-length bounded such that EPS = 0.0 at the root of the tree and EPS = around 100 at the leaf nodes. Nine clades in the tree are shown in the inner circular band. Fourteen clusters (with GG labels in red) from Fig. 1 are shown in the middle band for comparison with the 9 clades of the tree. The outer band shows the regional classification used in the SGDP. The blanks on the circular bands are for those individuals with no SDGP regional assignment (outer band), not clustered among GG0-GG13 (middle band), or not in any of the 9 clades (inner band). A corresponding linear topological tree with the names of EGs is shown in Supplementary Fig. S1.

a. As was observed in Fig. 1, all African GGs (GG0 – GG4) emerged sequentially in multiple steps, but all non-African GGs (GG5 – GG13) emerged in a burst within one giant clade nested by 4 large clades and a few small clades plus isolated branches;
b. We can identify 9 clades (inner circular bands), 8 of which are closely related to 8 out of the 14 GGs defined by the PCA clustering (middle circular band), and
c. Most of the 8 clades can be assigned to various geographical or geological regions.

The NJ tree of the first SDGP publication (5), when compared to ours, has several differences in clading pattern, relative branch lengths and branching order within each region (Extended Data Fig. 4B of reference *5*). These differences as well as those of clustering differences mentioned in previous section (**Clustering by PCA)** are ultimately due to the different ways the two methods describe the individual genomic variations, i.e., isolated SNVs in reference 5 vs. contextual SNVs in this study.

### Combining the clustering pattern in PCA, the clading pattern of NJ tree, and the clustering pattern in geographical map

Our GG classification is not based on PCA clustering alone. It is based on the cross-checking of the clustering pattern of individuals in PCA (in genomic divergence space), the clading pattern in NJ tree (in genomic divergence space), and the clustering pattern in geographical map. Fig 2 shows that 9 out of 14 GGs agree approximately between the PCA clustering and the clading in NJ tree (see the middle color band and inner color band, respectively). Among the remaining five GGs, four are based on the degree of agreement between the PCA clusters and much loose clusters in the current geographical locations of individuals (shown in the upper right inset of Fig. 1). For GG3, all three disagree in various extent. So we arbitrarily lumped together into one and called GG3 with the hope that, once more data become available in future, we may be able to subdivide into GG3a, GG3b, etc.

### Order of emergence of the founders of GGs on “Evolutionary Progression Scale (EPS)”

The order of emergence of all the extant GGs can be inferred from Fig. 1 based on the nearest-neighbor relationship among the centers of each GGs (see also Supplementary Fig. S2A and Table S3). Furthermore, the point on EPS (see ***“Evolutionary Progression Scale”*** in Supplementary Materials) at which the “founder(s)” of each GG emerged can be derived from Fig. 2 under the following processes: We start with two assumptions: (a) from the point of view of genomic information, the progression of evolution can be considered as the process of increasing divergence of genomic sequences; (b) the “founder(s)” of an extant GG can be considered as a selected subpopulation from the population of an internal node at a specific point on EPS (see the radial line scaled zero to 100 in Fig 2), and, then, the founders diversify and migrate to generate all the extant EG members of the GG. The genomic divergence, which corresponds to the cumulative branch-length, is set to EPS = 0 at the origin of our tree, and EPS = around 100 for the leaf-nodes of all extant individuals (see Fig. 2 and ***Evolutionary Progression Scale*** in Supplementary Materials). For example, the founder(s) of the first GG (GG0) emerges from the first internal node located at EPS = about 15.2 in Fig. 2.

Finally, we take the identification of each GG from Fig. 1 and its nearest-neighbor relationship from Supplementary Table S3, and combine with the EPS value for the corresponding internal node from our tree to estimate the order of the emergence of each GG *and* the point on the EPS of the emergence of the founder(s) of the GG in two spaces: one on PCA space (Supplementary Fig. S2A) and the other on a world map (Large circles in Supplementary Fig. S2B). Such combination suggests that:

a. All extant African GGs (GG0 – GG4) emerged *sequentially*, and their founders emerged between EPS of 15.2 and 29.4, but any presumably extinct earlier GGs must have emerged during the period corresponding to EPS between 0.0 and 15;
b. The first extant GG (GG0) consists of two EGs of Khomani_San and Jo_hoan (see Fig. 1 for the numbering of the GGs and Supplementary Table S1 for the names of EGs in each GG) currently residing in the southern tip of Africa, and the last African GG (GG4) is composed of the EGs of Mozabite and Saharawi in Northern Africa at the end of the African supergroup in Fig. 1;
c. The founders of the non-African GGs emerged in a *“*burst*”* into several lineages of the GGs during a period corresponding to a narrow range of EPS of 33.5 – 39.5 (see Fig. 2 and Supplementary Fig. S2A and B). They all emerged from GG5 (the first non-African GG nearest to the last African GG (GG4) in Northern Africa). GG5 consists of the EGs of Bedouin B, Druze, Iraqi Jew, Jordanian, Palestinian, Samaritan, and Yemenite Jew in the Middle East;
d. The founders of GG12 and GG13 emerged most recently at EPS of 39.5.
e. The EGs of a newly emerged GG are often found in a large region bound by great geolocal barriers different from those of previous GG, suggesting that some members of the previous GG may have migrated through the great geologcal barriers.

### Part II. Average genotype identity between two individuals

The genome-based grouping discussed above has been derived based on the “genomic divergent distance” between two FFPs of *contextually-linked* SNVs in each of all pairs of individuals. Although the distance is precisely defined from the Information-Theoretic viewpoint it is not obvious what the distance between the two FFPs corresponds to physically in terms of two respective whole genome contents. Furthermore, although 1KGP study (4) showed that the average SNV difference between the reference genome of European ancestry and each individual’s genome of different population groups vary from 3.54 million (M) to 4.32M SNP loci, corresponding to 4.2% to 5.1% of total SNP loci, it is not certain whether similar variations will be observed between two individuals among *all* study populations (not comparing each to the one Eurocentric reference genome). To get a more intuitive understanding, we ask alternative questions: (a) What is the average percent identity in terms of whole genome content between two individuals within one GG vs. from two different GGs? (b) Are the percent identity unique to each GG? and (c) What are the implications of the answers to the above questions?

For this portion of our study, genomic identity has been quantified in two steps: First, we quantify the genotype identity of all SNP loci within each pair of two individuals in each group as well as between two different groups, using the genomic data of the larger population size (the 1KGP data) and of the broader diversity in ethnicity (the SGDP data). Then, we combine the results to generalize and extrapolate to estimate an approximate average magnitude for the genomic identity between two individuals as a percent of the most recent *whole* human genome sequence with no gaps (about 3.06 billion (B) base-pairs (*15*). These steps are taken under a few approximating and simplifying assumptions described below:

There are many types of mutational events that result in individual genomic variations, such as single nucleotide substitutions, short Indels, large deletions/insertions, inversions and others. Of these, about 99.7% are due to SNPs, and the rest of variational events are one or more order-of-magnitude rarer than SNPs (*4*). Therefore, under the first approximation that SNPs account for the overwhelming portion of all mutational events, we ignore all other mutational events, which are very rare and difficult to quantify and compare. Thus, we estimate the extent of the individual SNV *identity* by counting identical genotypes between the genomes of two individuals at all available SNP loci, then we extrapolate (re-scale) the SNV identity among all SNP loci to that for a whole-genome length. This process was repeated for both data sets of the 1KGP (4) and the SGDP (5). We estimate the average percent-identity in four different ways, the results of which are summarized below. The detailed calculations are described in Supplementary Materials under ***Four ways of estimating the average identity of whole genomes between two individuals among the world’s ethnic groups***.

a. The average genomic identity between the human reference haploid genome, GRCh37 (*16*), was compared with each of all individual genomes from 26 “population groups (PGs)” of the 1KGP. Interestingly, the genotypes of 95.6%, on average, of all SNP loci (84.7M) between the reference genome and each individual is identical. Extrapolating or re-scaling this number to whole-genome length, 99.86% of the whole genome of each individual have identical genotypes as those of the reference genome under the simplifying approximations mentioned above.
b. Furthermore, the average genomic identity between two individuals (*excluding* the human reference genome) among *all* members of the 1KGP population is also found to be about 95.08% of the total SNP loci (Table 1A and Figs. 3A1 and 3A2), which corresponds to 99.87% of the whole genome of the recently updated human reference genome length (17, 18).
c. Similar to the results with 1KGP population above, the average genotype identity between two individuals among all members of 164 ethnic groups of the SGDP is 90.39% of a total of 34.4M SNP loci (Table 1B and Figs. 3B1 and 3B2), which corresponds to about 99.84%, on average, of whole genome length of SGDP.
d. Extrapolating these consistently high estimates for genome identity between two individuals in the 1KGP and the SGDP samples to the latest values of 111M for the whole genome SNP loci (18) and of 3.1B base-pairs for whole complete human genome length (15), we arrive at a “generalized” and conservative estimation of 99.8% as the genotype identity between two individuals regardless of the types of categorization by the “population groups” in the 1KGP or GGs of the ethnic groups of the SGDP.

**Figure 3.**
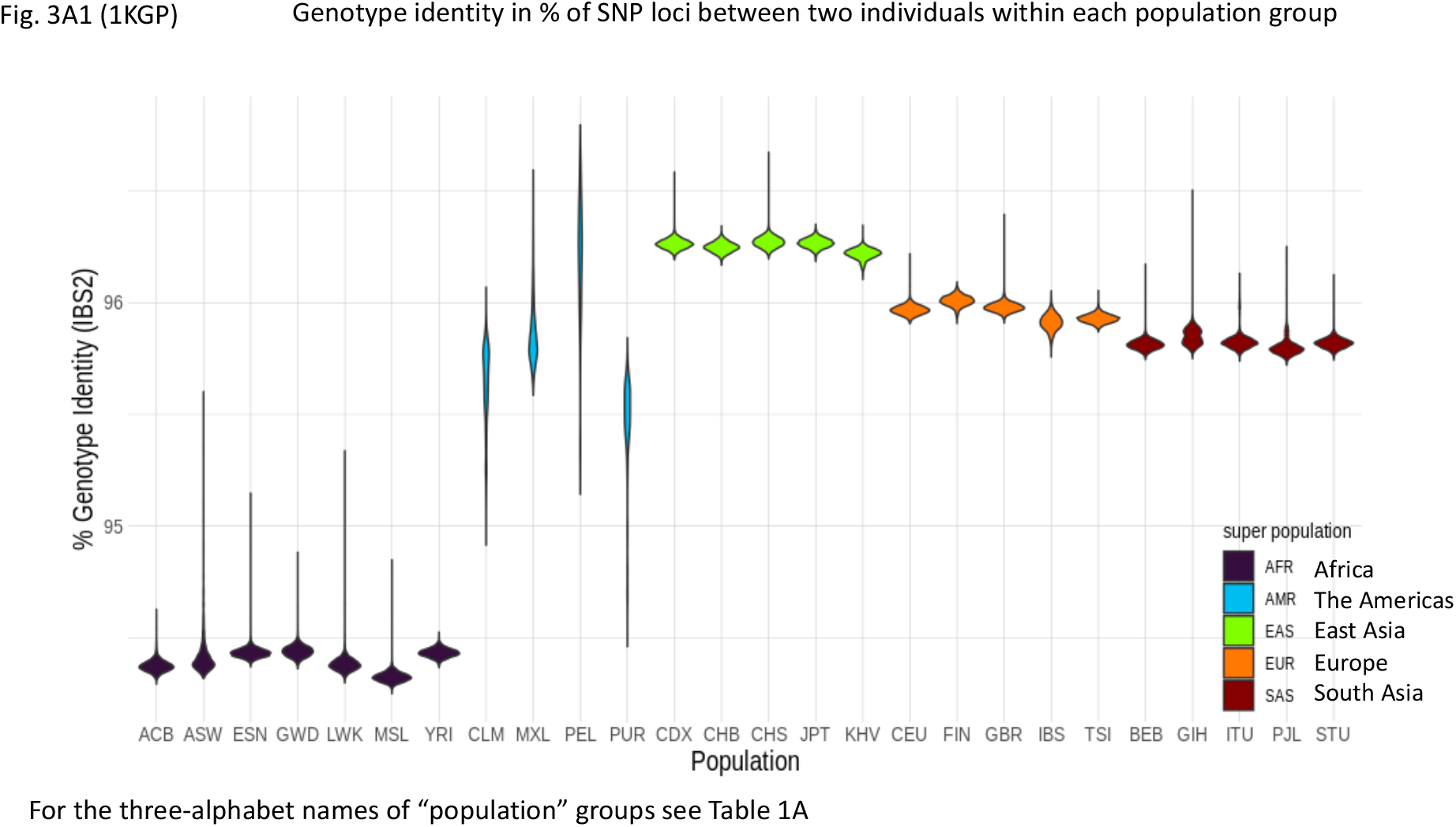
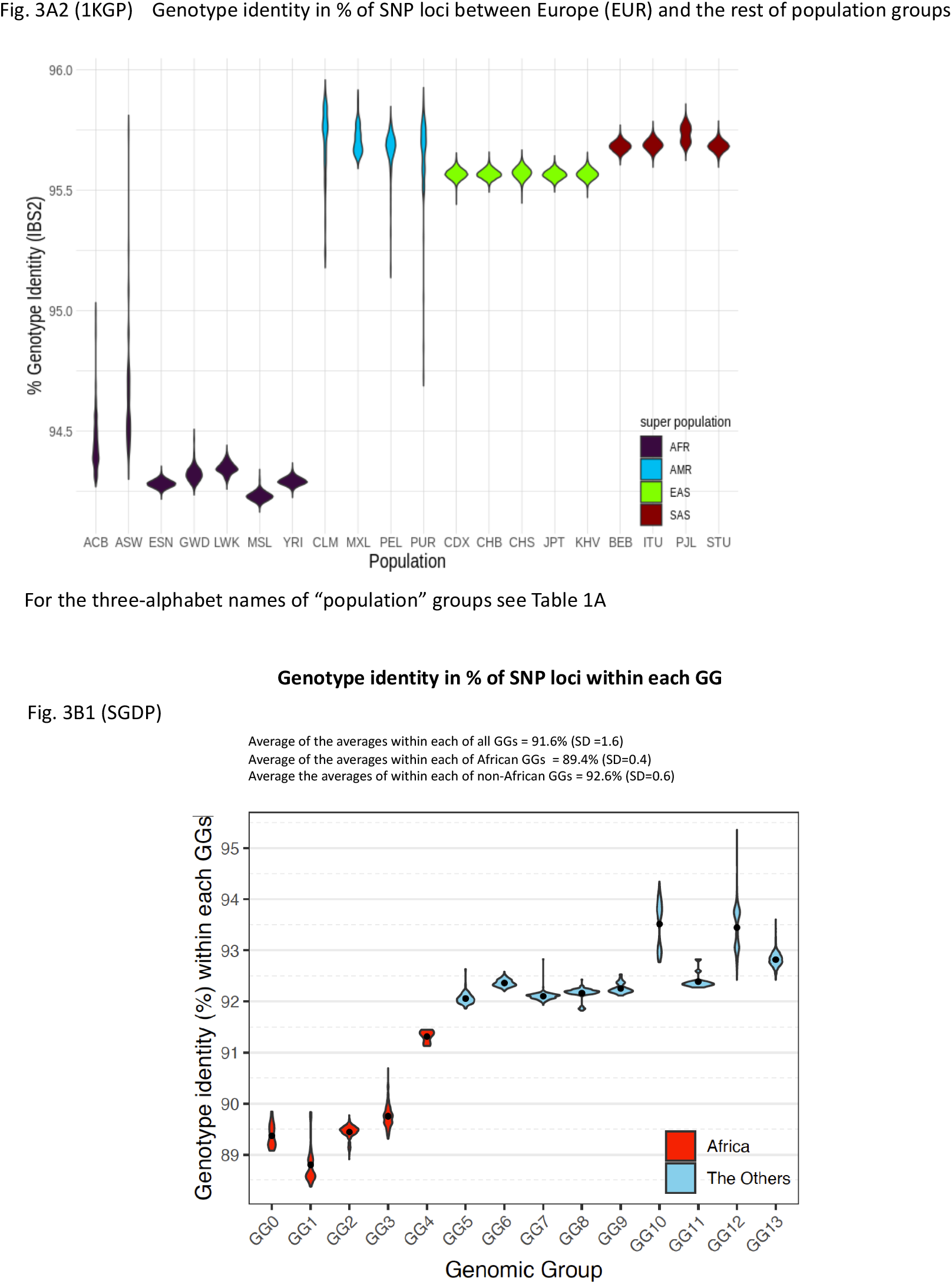
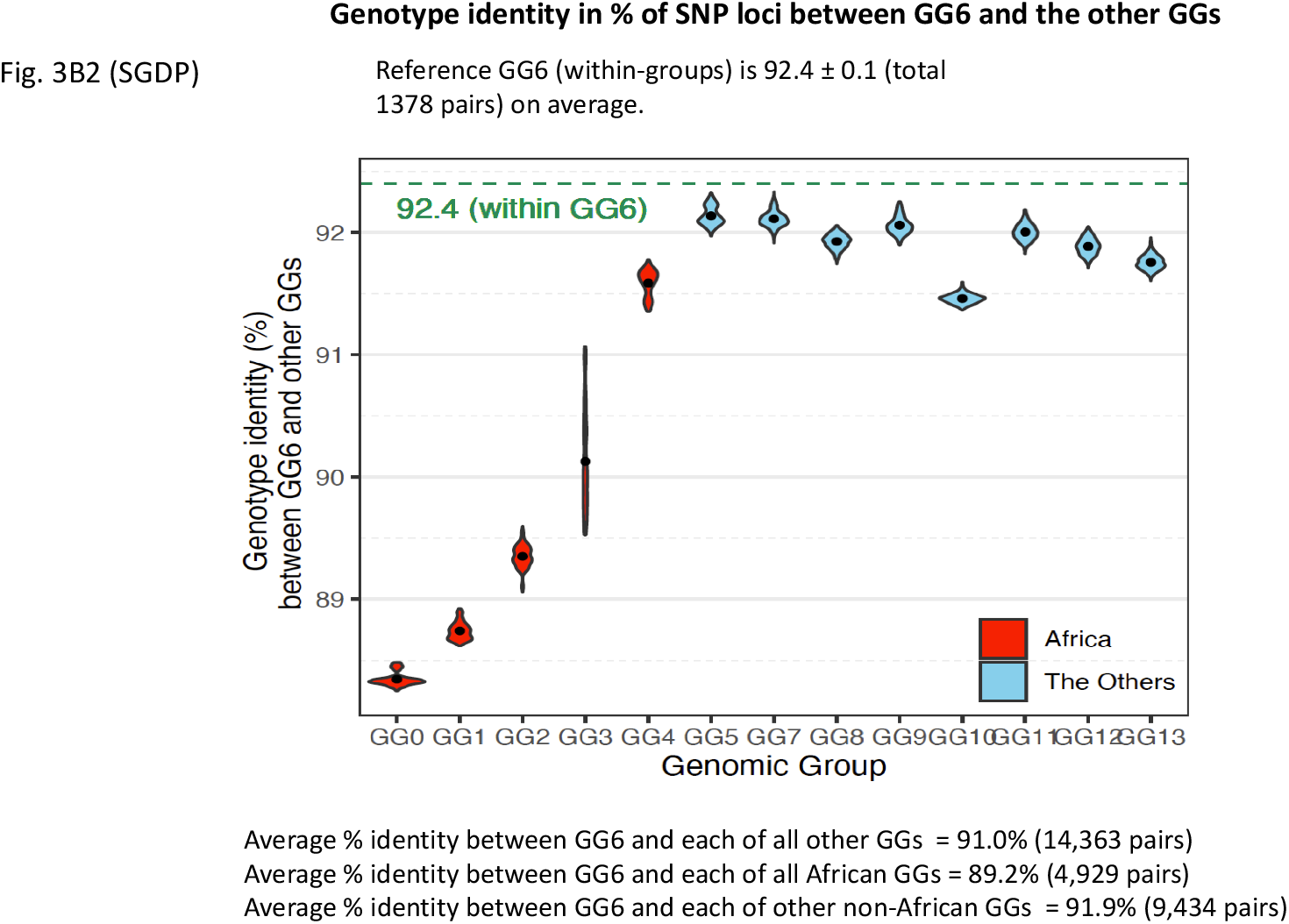
“Violin plots” of the genotype identity for all SNP loci. **(A1)** Violin plot of the % genotype identity for pair-wise SNVs among all SNP loci *within* each of 26 “population groups” of the 1KGP samples. The long streak for the distribution of % identity within each of 4 groups of the Americans reflect loose categorization by the population grouping of the American continental group in the 1KGP samples. The extent of the “streaking” is correlated to the standard deviation (SD) of each average value, which, in turn, is related to the “looseness” of each cluster (see Table 1A). The average % identities within African PGs (about 94.4%) is slightly lower than those of non-African PGs (about 95.9%), revealing that (a) African PGs and non-African PGs form two separate super groups, and (b) the African PGs have slightly more diverse genomes than those among the members of non-African PGs. **(A2)** Violin plot of the % genotype identity of pair-wise SNVs among all SNP loci *between* the European group (consisting of the PGs of CEU, FIN, GBR, IBS, and TSI) and the rest. The % identity between two different PGs are slightly lower than those within the same PG. The plots for those between each of non-European groups and the rest look similar, and not shown. (**B1**) “Violin plot” of the % identity of pair-wise SNVs among all SNP loci *within* each GG of the SGDP samples. The average of the averages of % identity within each of all GGs equals 91.6%, that within African GGs is 89.4%, and that within non-African GGs is 92.6%, revealing that (a) as in Fig. 3A1 with the 1KGP samples above, African GGs and non-African GGs of the SGDP samples form two separate super groups, and (b) the African GGs have slightly lower % genotype identity, i.e., slightly more diverse genomes than those among the members of non-African GGs within each GG. (**B2**) The % identity of pair-wise SNVs between GG6 and the other GGs. Plots for those *between* each of non-European GGs and the rest look similar, and not shown. The % identity between two different GGs are slightly lower than those within the same GG. The long streak for the distribution of % identity between GG6 and GG3 is because GG3 is not a tight cluster but a diffuse distribution of the individuals from several loose sub-clusters.

### Summaries and Prospects

At the level of the population of all 164 ethnic groups in the SGDP, we show the feasibility of objectively categorizing the ethnic population into approximately 14 exclusively genome-based groups (GGs) without any consideration of self-declared or subjectively perceived categorization, such as race, ancestry or ethnicity. Such genome-based categorization should provide a scientific footing for demographic studies of various areas such as:

a. Estimating or predicting the role of the inherited genomic components that contribute to certain phenotypes specific to a given GG in the health-related fields, such as epidemiology, disease diagnosis, disease susceptibility, and clinical practice in predicting, for example, drug or therapy responses;
b. Training and testing of various computational algorithms using Information Technology, e.g., Machine Learning and Artificial Intelligence in the fields mentioned above, where the development of such algorithms depend on having objectively definable demographic categorizations (called “labels” in this field) of the genomic input data.
c. Supplementing current non-genomic characteristics with our genomic characteristics to improve human categorization in general.
d. Suggesting the need for additional genomic diversity data for the ethnic groups especially in Northern Africa, Nordic Europe, North and Central Asia, and the Americas, for which currently existing data are very sparse. Such data may lead to discovering additional GGs.

At an individual level, taking all four estimates derived above together we make a simplified and conservative approximation that the genomic identity is, on average, about 99.8% of whole genome (ranges of the four estimates above: 99.82% - 99.87%) between two individuals regardless of categorization, i.e., the remaining 0.2% accounts for non-identical genotypes, which amount to about 6M genomic loci. This approximation is also under the final assumption that the degree of the genomic diversity of the ethnic groups in this study represents, approximately, the genomic diversity of the extant human species. Possible implications derived from the observation of the uniformly very high genomic identity between two individuals regardless of categorization are discussed below in last three sections of **Implications and Discussions**.

### Implications and Discussions

The genome-based grouping pattern and individual genomic identity described in Results are the *average* features of the grouping and quantifications for the *majority* of the extant ethnic and population groups at the time of this study, when the field continues moving toward obtaining higher quality (*18*) and completeness (*15*) of the reference human genome sequence.

Not discussed in the Results are: (a) those features contributed by the “outliers” with rare degrees of genomic identity/difference between two individuals located in the long “tails” of the main bodies of the “violins” in Fig. 3, and (b) those contributed by other inherited but non-genomic elements, such as epigenomic chemical modifications of genome, non-genomic but inherited nucleic acids, microbiome, and others (*19*), for which no comprehensive data are available, at present, to make quantitative estimates of their contributions to a genome-based categorization. With these caveats we describe the implications of the Results with discussions below.

### “Burst” of all extant non-African ethnic groups from the Middle Eastern GG

**O**ur observations in part I of the Results above imply that: (a) the most recent ancestors of the extant ethnic groups in this study emerged sequentially from Southern Africa (GG0), and (b) all founders of the non-African ethnic groups emerged in a “burst” from the Middle Eastern genomic group (GG5), which emerged after the two Northern African ethnic groups of Mozabite and Saharawi (in GG4) in Northern Africa emerged. What event or events may have caused such burst?

### About 99.8% genomic identity for individuals and 96.4% for all populations

Under the simplifying assumptions mentioned in part II of the Results above and at the level of order-of-magnitude, we find that the contents of the whole genomes of any two individuals in this study populations are very similar (about 99.8 % identical on average (with a narrow range of 99.82% - 99.87%) between two individuals not only within a given GG or PG but also from two different GGs or PGs. On the other hand, the genomic identity at all SNP loci among *all* PGs in 1KGP is about 96.4% of the whole genome of 3.1B base-pairs (see section (d) of ***Four ways of estimating the average identity of whole genomes between two individuals among the world’s ethnic groups*** in Supplementary Materials). This observation infers that the loci with different genotypes between two individuals in one pair is different, in general, from those in a different pair, although the magnitude of the difference is about the same for both pairs. Thus, the union of the loci with identical genotypes among *all* pairs is significantly lower than the number of loci with identical genotypes *between* two individuals, i.e., all members of the study population as a single group has 96.4% identical genome, but, at an individual level, 99.of whole genome is identical between two individuals, in general. One more detail: the average % SNP identity between two individuals among African GGs or PGs is about 1% lower than that among non-African GGs or PGs, but when extrapolated to whole genome the 1% difference of SNPs corresponds to less than 0.03% of whole genome length. This is also true when we compare the SNP identity between one individual from African groups and the other from non-African groups (see Tables 1A and 1B). Thus, the genomic divergence among the members of African GGs or PGs are always slightly higher than those of non-African GGs or PGs, as widely accepted in the field.

### Inherited “Passive” genomic information

Our observation of almost uniform degree of very high genomic identity (99.8%) indicates that any two extant individuals have inherited an *almost complete* set of identical genomic information, despite apparently complex phenotypic differences. This apparent “inconsistency” implies that the information in the inherited whole genome may be a near-complete set of *“passive” information (or potentials*) that are differentially activated by a combination of the diverse non-genomic (environmental) factors and a very small fraction (0.2% of whole genome) of an individual’s inherited genome (see next paragraph, and **Indirect implication on a molecular scenario for converting environmental diversity to genomic divergence** in ***Supplementary Note 2*** in Supplementary Materials) for basic survival and reproduction of the individuals. For example, a frozen embryo does not have life, although it has the complete set of genomic information and essential proteins and other chemicals to start life, until certain environmental signals, such as heat and essential nutrients, are received to start its life. In this scenario the environmental signals, be they from the “biological environment” (in-utero, microbiome, ecology, food, family, society, culture, faith, ideology, lifestyles, etc.) or “non-biological environment” (heat, radiation, geology, climate, atmosphere, etc.), can be considered as a part of critical “activating” agents of the “passive” genomic information.

### Environmentally-selected genomic variants

Although most of each GG form tight genomic clusters, the members of each GG are found spread broadly within one of about 11 geological/geographical regions (S. Africa, Mid Africa, N. Africa, Middle East, Europe, Central Asia, S. Asia, Oceania, North Asia, the Americas plus NE Asia, and E. plus SE. Asia) as shown in Fig. 1 and Supplementary Table S2. Many of the regions are definable by various major geological barriers such as high mountain ranges, large body of water or desert, etc. For example, the members in GG6 are widely spread in Europe, but segregated from those of other non-African GGs by the Ural mountain range and Caspian sea; GG7 and GG8 (both in S. Asia) are separated by a large geological barrier, Thar desert; most members of GG12 are in the North and South America; and the members of GG13 (in E. and South East Asia) are segregated from those of the rest of Asia by Himalaya mountains on the South, Tianshan mountains, Alta mountains and the Gobi desert on the North. Thus, interestingly, the categorization of GGs, which are exclusively genome-based, appears to correlate with non-biological environment bordered by major geological barriers. This observation implies or suggests that even the genome-based categorization of a GG may have been strongly influenced by the environmental selection of a unique set of genomic sub-variants (from a diverse genomic variant population accumulated during a long environmental exposure specific to the survival advantage of the GG) on an evolutionary time scale.

### Environment-based vs. genome-based categorization

The fact that there is no one-to-one correspondence between the 14 *explicitly* genome-based GGs and the 164 EGs (as designated based on “ethnicity” in the SGDP) or the 5 “racial groups” (Black or African Americans, White, Asian, Native Hawaiian or Other Pacific Islander, and American Indian or Alaska Native, as defined in US 2020 census based on skin color and geographical regions) (see Supplementary Table S3) indicates that the biologically inherited genomic diversity does not play significant role in the categorization by the current ethnicity or race. From the genomic view point, the overwhelming portion of the inherited genomes, which is about 99.8% identical between two individuals, do not play a very influential role in the categorization of ancestry, ethnicity or race. Furthermore, even the genome-based categorization of GGs is not only derived from a very small fraction (0.2%) of the whole genome, but also may have originated from the environmental selection of the genomic variations of the 0.2% (see previous paragraph). Thus, our results imply that both ethnicity and race are non-genomic, i.e. *environmental*, categorization, be they biological and/or non-biological.

## Data Availability

All genomic data used in this study have been released and are publicly available from:

a. the Simons Genome Diversity Project (https://www.simonsfoundation.org/simons-genome-diversity-project/ ; https://reichdata.hms.harvard.edu/pub/datasets/sgdp/) and
b. the 1000 Genomes Project (https://www.internationalgenome.org/data ; https://www.internationalgenome.org/data ; http://ftp.1000genomes.ebi.ac.uk/vol1/ftp/release/20130502). For this study, The SNP data from The Simons Genome Diversity Project database (*5, 6*) were last accessed in July of 2020, and those from The One Thousand Genome Project database (*4, 18*) were last accessed in June of 2020.

No additional new genomic data have been generated from this study.

## Acknowledgements

We are grateful to the Simons Genome Diversity Project and the 1000 Genomes Project for making their SNP data available publicly. We thank Prof. Rajiv McCoy of Johns Hopkins University for their updated information on the first complete human genome sequence of CHM13 (*15*) and Dr. Michael Zody of New York Genome Center, NY, USA on updated information about human reference genome GRCh38 (*18*). SHK acknowledges having appointments as Visiting Professorships at Yonsei University, Korea Advanced Institute of Science and Technology, and Incheon National University in South Korea during the beginning part of manuscript preparation.

## Author contributions

Conceptual design of the study and speculative interpretations and implications of the results by SHK; all computational work including algorithm design, programming and execution for this study was done mostly by BJK and Neighbor Joining tree construction by JJC; discussion and interpretation of computational results by BJK, JJC and SHK; manuscript preparation by SHK with extensive discussions with BJK and JJC; figures designed by BJK, JJC and SHK.

## Computer code availability

The codes to parse VCF files, determine/visualize PCA and display geological locations are available at https://github.com/UCB-KIMLAB/peopling, and the programs to generate JSD distance matrix are available at https://github.com/jaejinchoi/FFP, using the version 2v.3.1.

