## Supplementary materials for "On whole-genome demography of world’s ethnic groups and individual genomic identity"

I. **Data source and Methods**

***SNV data source:*** For the availability of all genomic data used in this study see **Data Availability** in the main text. All data have been released and publicly available from

The Simons Genome Diversity Project (*5, 6*) and The 1000 Genomes Project (*4, 18*). The Simons Genome Diversity Project database were last accessed in July of 2020, and those from The 1000 Genomes Project database were last accessed in June of 2020.

***Feature Frequency Profile (FFP):*** For the purpose of introducing the concept of the contextual SNV (c-SNV), each individual’s whole genome SNVs are represented by a vector, FFP (*22*), consisting of multiple millions of unique c-SNVs, as genomic characters, and their respective frequencies, as character states. Each c-SNV is a short string of linked SNVs of a given length (l), which has l -1 overlapping sequence of its neighbor c-SNVs. Since FFP is a collection of all overlapping c-SNVs and respective frequencies, it contains all information needed to reconstruct the original ordered SNVs, thus, an FFP of c-SNVs is a convenient vector to represent all SNVs of an individual’s whole genome for mathematical comparisons and manipulations.

***Optimal feature length of c-SNVs:*** One of the critical parameters in FFP method is to predict the optimal length of c-SNVs, but there is no *a priori* method to find it. We, therefore, take an empirical approach to find the optimal Feature length by constructing a series of FFP-based trees using c-SNVs of incrementally longer lengths and choosing the length that produces the most topologically stable tree when compared with its neighbor trees. The results for this study are shown in Supplemental Fig. S3.

***FFP-based PCA****:* The input for the classical PCA used in this study is the pair-wise “distance” matrix, where the distance is calculated by pair-wise Jensen-Shannon divergence between two leaf nodes represented by two FFPs (see **Jensen-Shannon divergence and “Evolutionary distance matrix”** below)

***FFP-based rooted Neighbor-Joining (NJ) tree*:** The neighbor-joining trees (13) in this study were built using BIONJ (*14*). Two unique differences in this study are: (a) as in PCA above, the starting point is the pair-wise “distance” matrix, where the distance is calculated by pair-wise JS divergence between two leaf nodes represented by two FFPs (see **Jensen-Shannon divergence and “Evolutionary distance matrix”** below); and (b) the tree is rooted by an artificial out-group (see **Out-group** below). Since each branch-length is proportional to the genomic divergence as a result of evolution, the cumulative branch-length from the root to a given internal node represents the extent of evolutionary progression of the node.

***Jensen-Shannon (JS) divergence and “Evolutionary distance matrix”*** ***:*** The “genomic distance” between two FFPs for a given l, represented by two frequency vectors P and Q for all of the equivalent c-SNVs, are calculated by the JS divergence (*23*):

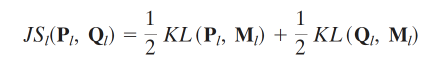

Where M = (P + Q)/2 and *KL* is the Kullback-Leidler (KL) divergence:

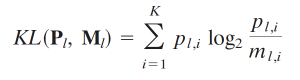

For vector Q, replace P and p by Q and q in the above equation, where p and q are the components of P and Q vectors.

*K* is the union of all unique c-SNVs in P and Q, and runs usually in millions. JS divergence has useful properties for our study: They are symmetric and bound between 0 and 1 for two identical FFPs and two completely different FFPs, respectively; they reflect the extent of genomic divergence as evolution progresses. The collection of all JS divergence provides a matrix, which we take as the “evolutionary distance matrix”, and use as the starting input for PCA as well as for the construction of neighbor-joining trees.

***Out-group:*** Since our FFP method does not require multiple sequence alignment of the linked SNVs, we construct the SNV sequence of an artificial outgroup member by taking one actual SNV sequence and shuffling their sequences. Such synthetic and artificial SNV sequences would have the same length and composition as the actual SNV sequence without the contexts of the true SNVs. The rooting based on such synthetic SNVs as the outgroup allows us to identify the first emerging GG as two EGs of Khomani_San and Jo_hoan of Africa.

***“Evolutionary progression scale”:*** In Information Theory, the Jensen-Shannon (JS) divergence, bound between zero and one, is commonly used as a measure of the dissimilarity between two probability distributions of linear informational features. The FFP as the distribution of frequencies (character states) of unique c-SNVs (genomic characters) is such a probability distribution. Thus, a JS divergence of two FFPs, used as a measure of the genomic-divergence distance between two FFPs of c-SNVs, is also bound between 0 and 1, corresponding to the JS divergence value between two identical FFPs, representing two organisms with the identical genome, and two completely different FFPs, representing two organisms with completely different genomes, respectively. In addition, since the genomic divergence is defined mathematically similar to physical entropy, which increases on time progression, the JS divergence will have the same direction and magnitude of evolutionary progression, thus “evolutionary progression scale”. Any whole c-SNV “dissimilarity” between two individuals accumulated during the evolution from their common ancestor can be considered as the sum of the two divergences, each expressed as cumulative branch-length from the common ancestor to respective leaf node as caused by changes of, ultimately, genomic sequences due to all types of mutations, such as point substitutions, indels, inversion, recombination, loss/gain of genes, etc. as well as other unknown mechanisms, and they will bring JS divergence somewhere between 0.0 and 1.0 depending on the degree of the sequence divergences from the common ancestor.

In this study the collection of the JS divergences for all pairs of the study individuals plus 2 out-group members (see “**Outgroup”** in Materials and Methods) constitutes the “distance matrix” for BIONJ (*13, 14*). Since all the branch-lengths are derived from the JS divergence values, the cumulative branch-length of an internal node corresponds to “cumulative genomic divergence (CGD)” of the node along the presumed evolutionary lineage. Thus, it can be considered as the point of evolutionary stage reached by the node on an “evolutionary progression scale”. For convenience of assigning the nodes on the progression scale, CGDs are scaled such that the CGD value at the root node of the tree is set to zero and the leaf nodes of the individuals to 100, on average, corresponding to the fully evolved genomic states of the individuals at sampling time, which we define as the end point of the “evolutionary progression scale” for the extant individual. (see Fig. 2, where the cumulative branch-lengths are multiplied by 200 for easy visibility as CGDs, which are used to represent “Evolutionary Progression Scale (EPS)”)*.*

***Four ways of estimating the average identity of whole genomes between two individuals among world’s ethnic groups:***

**(a). Average genomic identity between one human reference genome and each of all individual genomes from 26 “population groups (PGs)” of the 1KGP**: The first hint of the extent of the genotype difference between two individuals came from the 1KGP (*4*), where the genotypes of whole genome SNP loci of each of 2,504 individuals from 26 PGs of the 1KGP were compared with those of a single reference haploid human genome, GRCh37 (*16*), which is not of the genome of one individual, but a synthetic hybrid of multiple individuals’ genomes of mostly European ancestries with only the ABO blood type of “O”. Comparing each of all individual genomes to this single reference haploid genome, the study found that, on average, only about 3.58 million (M) (for non-African individuals) to 4.31M (for African individuals) SNP loci out of a total of 84.7M SNP loci (for whole “callable” genome that excludes the regions for which genomic variant calls are difficult) have genotypes different from those of the reference genome (from Table 1 of reference 4). When averaged over the 5 regional populations (Africa, Americas, East Asia, Europe, and South Asia), 3.73M SNP loci (4.4% of total 84.7M SNP loci) show individual genotypes (SNVs) different from those of the reference genome; thus, 95.6% of SNP loci have identical genotypes, on average, between the reference genome and each of all study population. At this point we make the second approximation that the “ungapped” reference genome GRCh37, which has about 2.87 B basepairs (16), may be a reasonable proxy for the “genotype-callable” genome. Extrapolating the 95.6% of SNP loci identity within all SNP loci to the whole “ungapped” genome of the reference genome (which is about 92.5% of the total GRCh37 sequence length, but excludes the regions such as pericentromeric and subtelomeric regions, ampliconic gene arrays, and ribosomal DNA arrays (*15, 16*)), 99.86% of the ”ungapped” whole genome of each individual, on average, have identical genotypes as those of the reference genome under the two simplifying approximations.

**(b). Average genomic identity between two individuals among 26 PGs of the 1KGP:** However, the above result is biased toward European genomes, because the reference genome, to which each of all other individual genomes are compared, is constructed from multiple European-descendant genomes. To avoid potential effects of the bias we calculated, for the second estimation, the genotype identity of all-to-all pairs (3,103,786 pairs) from 2492 individuals of the 1KGP **(**after excluding the reference genome and filtering out 12 kinship related samples from 2504 samples) from the 26 PGs. Table 1A shows a summary of the results averaged for each of the 26 PGs of 5 continental groups in four different ways: The average of the SNV identity between the pairs of all individuals within each of all 26 PGs equals 95.54% (with a standard deviation (SD) of 0.05%); that between one from each of African PGs and the other from the rest of PGs is 94.35% (SD 0.11%); that between one from non-African PGs and the other from the rest, is 95.35% (SD 0.14%); and that between one from all groups and the other from the rest of all groups is 95.08% (SD 0.48%). Thus, the overall SNV genotype identity between two individuals is found to be about 95.08% (range: 94.26% - 95.39% per all PG), on average, of total SNP loci of 84.7 M. The narrow range of the % SNV identities indicates a very high genomic identity between two individuals regardless of whether they are from the same or different PGs. Extrapolating the % average SNV identity among all SNP loci to 2.87 B base-pairs of the whole length of the “ungapped” reference genome of GRCh37 (*16*), we get 99.87% (range: 99.83% - 99.86% per PG) as an overall genotype identity, on average, for the “ungapped” GRCh37. Almost identical % identity is found even when we use the recently updated human reference genome of GRCh38.p13, which shows a longer “total ungapped length” (*17*) and an increased number of total SNP loci (*18*)). For easy visualization of Table 1A, two examples are given: Fig. 3A1 shows the SNV % identities of the genotypes among all SNP loci between two individuals within each of the 26 PGs (blue column of Table 1A), and Fig. 3A2, as one example among many between-group comparisons, shows that between each of five EUR (European) populations and the rest of all other population groups. It is noticeable that the % genotype identity of SNP loci within as well as among seven African PGs is slightly lower (about 1 %), i.e., slightly more diverse, than those of non-African groups (Fig. 3A1), consistent with the segregation of all African GGs and all non-African GGs observed with PCA clustering in Fig. 1. However, this small difference, when extrapolated from SNP loci to whole genome, it becomes negligible.

Thus, under the approximating assumptions, all pairs of two individuals in the study population have identical genotypes to each other at 99.87%, on average, of their “ungapped” genomes.

**(c). Average genomic identity between two individuals among all members of 164 ethnic groups of the SGDP:** So far, the above two estimates are based on the 1KGP data that covers 26 PGs, which are much broadly defined geographical/ethnic population groups than the ethnic groups of the SGDP based on genetic, linguistic and cultural variations. Thus, for our third estimation, we examined the degree of genotype identity between two individuals of the study population of the SGDP database containing 164 ethnicity-based EGs, where the genomes were sequenced to an average coverage of 43-fold, which is much deeper than that of 1KGP. However, we cannot directly compare the SGDP study (*5*) with that of the 1KGP (*4*), because in the SGDP study the authors “retained” an average of 2.13B base-pairs per genome after filtering at “filter level 1”, and identified 34.4M SNP loci (in contrast to 84.7M SNP loci out of 2.87B “ungapped” length of the reference genome (*16*) in the 1KGP) also using the GRCh37 reference human genome. Thus, we compare all-to-all pairs (59,340 pairs) of the whole genome SNP genotypes from 345 individuals in the SGDP database within the “filtered” genomes. The results of comparison of all pairs of the SGDP individuals are summarized in Table 1B. On average, 90.39% (range: 88.31% – 91.13%, in the last yellow column of Table 1B) of 34.4M SNP loci have identical genotypes. Extrapolating the average SNV identity among all SNP loci of SGDP to 2.13B base-pairs, on average, of the “filtered” whole genome represented in the SGDP data indicates that 99.84% (range: 99.81% - 99.86% per GG) of the “filtered” genome has identical genotypes, on average, between two individuals. For easy visualization of Table 1B, two examples are given: Fig, 3B1 shows the % SNV identities of the genotypes among all SNP loci between two individuals within each given GG, and Fig. 3B2, as one example among many between-group comparisons, shows that between each of GG5 (European) populations and each of the rest of all other GGs groups. It is also noticeable, as with 1KGP population, that the % SNV identity within as well as among African GGs is slightly lower, i.e., slightly more diverse, than those of non-African GGs, consistent with the segregation of two super-groups of all African GGs and all non-African GGs observed with PCA clustering in Fig. 1.

Thus, under the approximating assumptions, all pairs of two individuals in this study population have, on average, 99.84% identical genotypes to each other in their “filtered” genomes.

**(d). “Generalized” average genomic identity of 99.8% between two individuals of world ethnic population:** Continuing advances in quality and quantity of whole genome sequencing techniques suggest that we can expect many more complete genome sequences with no gaps and no “unplaced bases”. Such advance has already been apparent with the recent *complete* telomere-to-telomere sequence of about 3.06B base-pairs (without Y chromosome) of CHM13 **(***15***),** which also identified 110M SNP loci for the population of the 1KGP. Similarly, an updated GRCh38.p13 sequence shows an “ungapped” length of about 2.95B base-pairs out of an estimated 3.10B total sequence length (*17*), which also identified 111M SNP loci (*18*) for the population of the 1KGP with respect to GRCh38.p13. These observations suggest that the total whole genome sequence lengths of CHM13 and GRCh38.p13 as well as the numbers of the total SNP loci for over 3,000 individuals of the study population of the1KGP may be reaching “asymptotic” values: The total genome length of human genome with no gaps of about 3.1B base-pairs and the total size of SNP loci of about 111M. These recent values also allow us to make a *conservative* estimation that the genomic identity at the level of all populations of the 1KGP is about 96.4% of whole complete genome, i.e., (3.1B - 111M) / 3.1B.

Thus, for our fourth estimate, we use the % SNP loci identity of 95.08% between two individuals (in section (b) above for the 1KGP sample), combine with the asymptotic values of 111M for the whole genome SNP loci and of 3.1B base-pairs for whole complete genome length, and arrive at the estimation of 99.82% as the % genotype identity between two individuals.

II. **Supplementary Notes**

***Supplementary Note 1:*** Some notable unexpected-observations in the PCA plot are listed below.

1. There are two Bantu Tswanas, but one belongs to GG1 and the other to GG2
2. There are 4 Punjabis, one of which is in GG7 and the other three in GG8
3. There are two Daurs: Daur2 in GG13 and Daur1, which has the highest missing SNVs, as an un-clustered individual near to singleton Hawaiian.
4. Maori belongs to GG8 in India although its current geographic location is in New Zealand.
5. Tlingit2 belongs to G9, but Tlingit1 is next to unclustered Saami pair sampled in N. Finnland.
6. Most of ethnic groups in Southeast Asia, including many islands, cluster with G13 or GG8.

***Supplementary Note 2*: Indirect implications on a molecular scenario for converting environmental diversity to genomic divergence**. Such a high genomic identity (99.8%) leaves only about 0.2% of the whole genome content to account for any genotype and phenotype differences between two individuals from within a given PG/GG or between two different PGs/GGs. In theory, it is possible that this 0.2% may be just random genotypes uniformly spread through all PGs/GGs. But that possibility is highly unlikely because such uniform noise would not produce 13+ genome-based distinct clusters of the study population as we have shown in Fig. 1. A more likely possibility is that this 0.2%, which amounts to about 6M genomic loci located mostly outside of the coding regions, may be non-randomly distributed genotypes and they participate in regulating, directly or indirectly, the transcription of a set of key genes in response to environmental signals, thus generating environment-dependent phenotype/trait variations. This possibility implies that this 0.2% of the whole genome genotype differences may play a role in connecting the environmental (non-genomic) signals, be they biological or non-biological, to the individual’s genomic variations of the 0.2% to produce phenotype diversity for the population. Then, some of the phenotypes are selected due to better adaptability to the given environment, and the genotypes of the selected individuals are inherited for the subsequent generations, thus, environmental selection becoming genomic selection such GGs. Furthermore, this bottlenecking step of the environmentally selected subpopulation with non-random sets of genotypes from the 0.2% results in forming the founder group of a specific GG, which appear to correlate with geological regions (see “**Interconnection between environment and individual’s genomic variations”** under **Direct Implications and Discussions)**

Above scenario assumes three types of events:

(a) “**Genomic diversification**” (including “genetic drift”) of the founder population of the all extant groups, resulting in the genomic variation of 0.2% between two individuals as the “base-line genomic noise” (for example, due to errors caused by the biological environment such as the errors during DNA replication or/and repair, or non-biological environment such as radiation and environmental mutagens), thus generating genomic diversity

(b) “**Phenotypic selection”** by one or more cycles of environmental (including “natural”) selections of the phenotype variants, which carry non-random set of genotypes from the 0.2%, better suited for the environment of each GG; and

(c) “**Genotypic selection**” by the sexual reproduction among the population of the phenotype-selected variants (thus, non-randomly selected genotype variants) in each GG.

Multiple cycles of the three types of events would ultimately form a particular GG with a set of distinct genomic variations at the level of about 0.2% of their whole genomes with apparent correlation to different geological regions.

III. **Supplementary Figures and Tables**

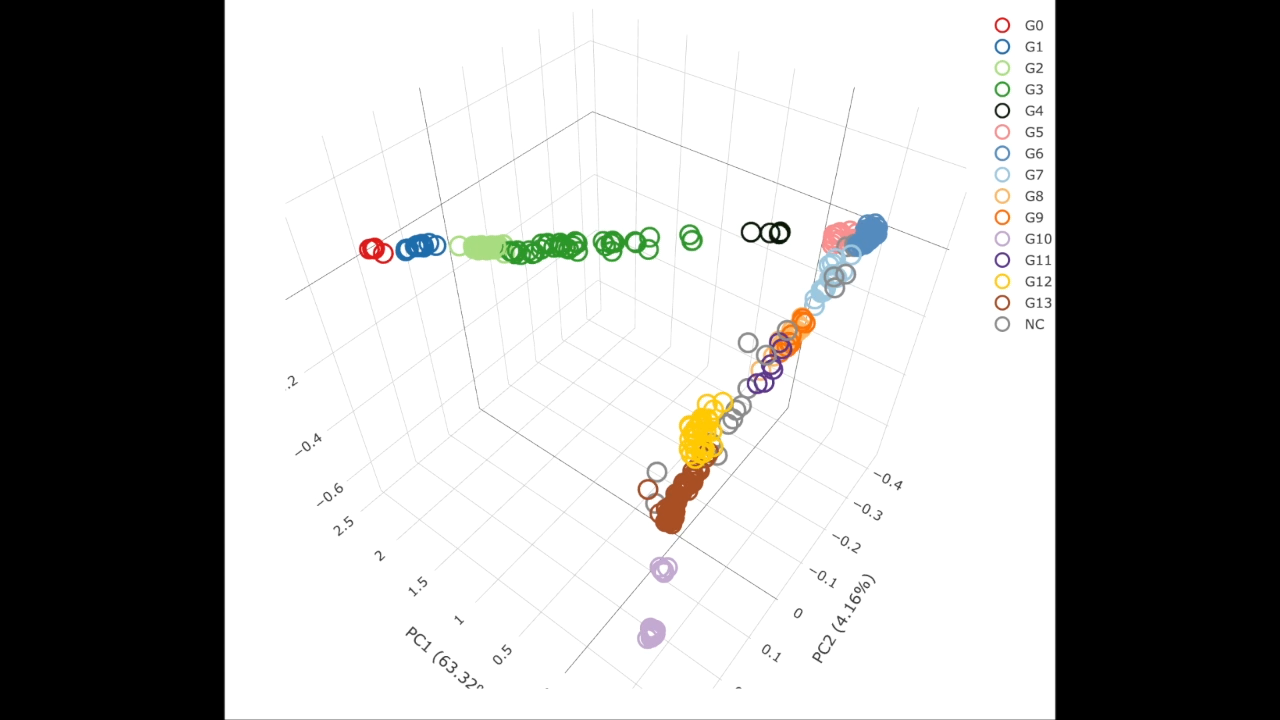

**Supplementary Fig. S1A. A video of the rotating view of the 3D PCA plot**

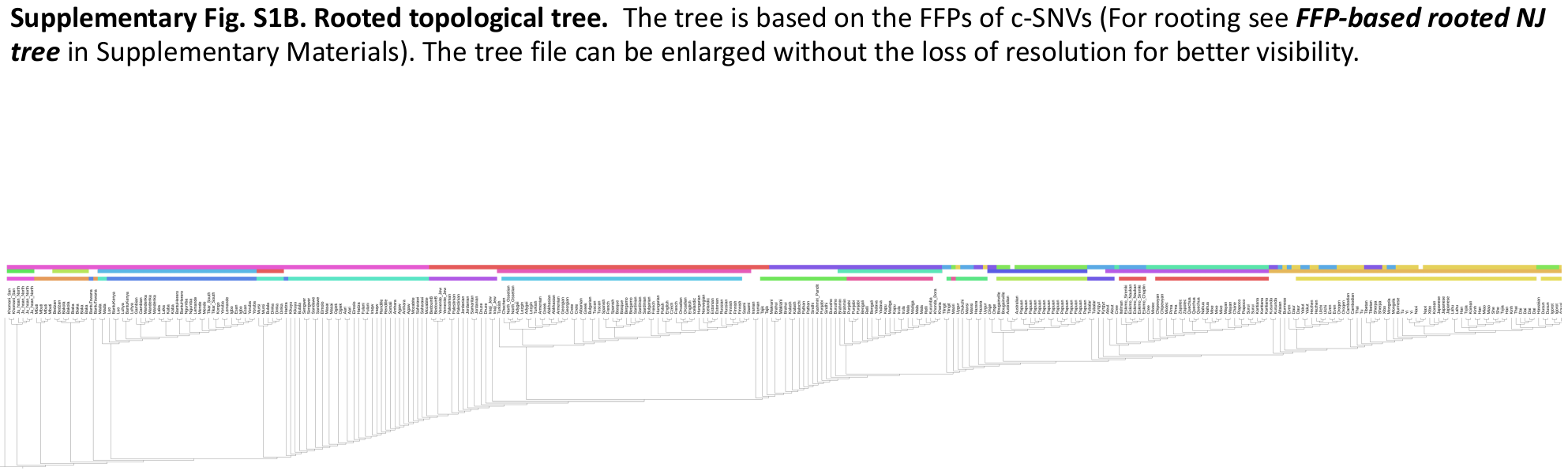

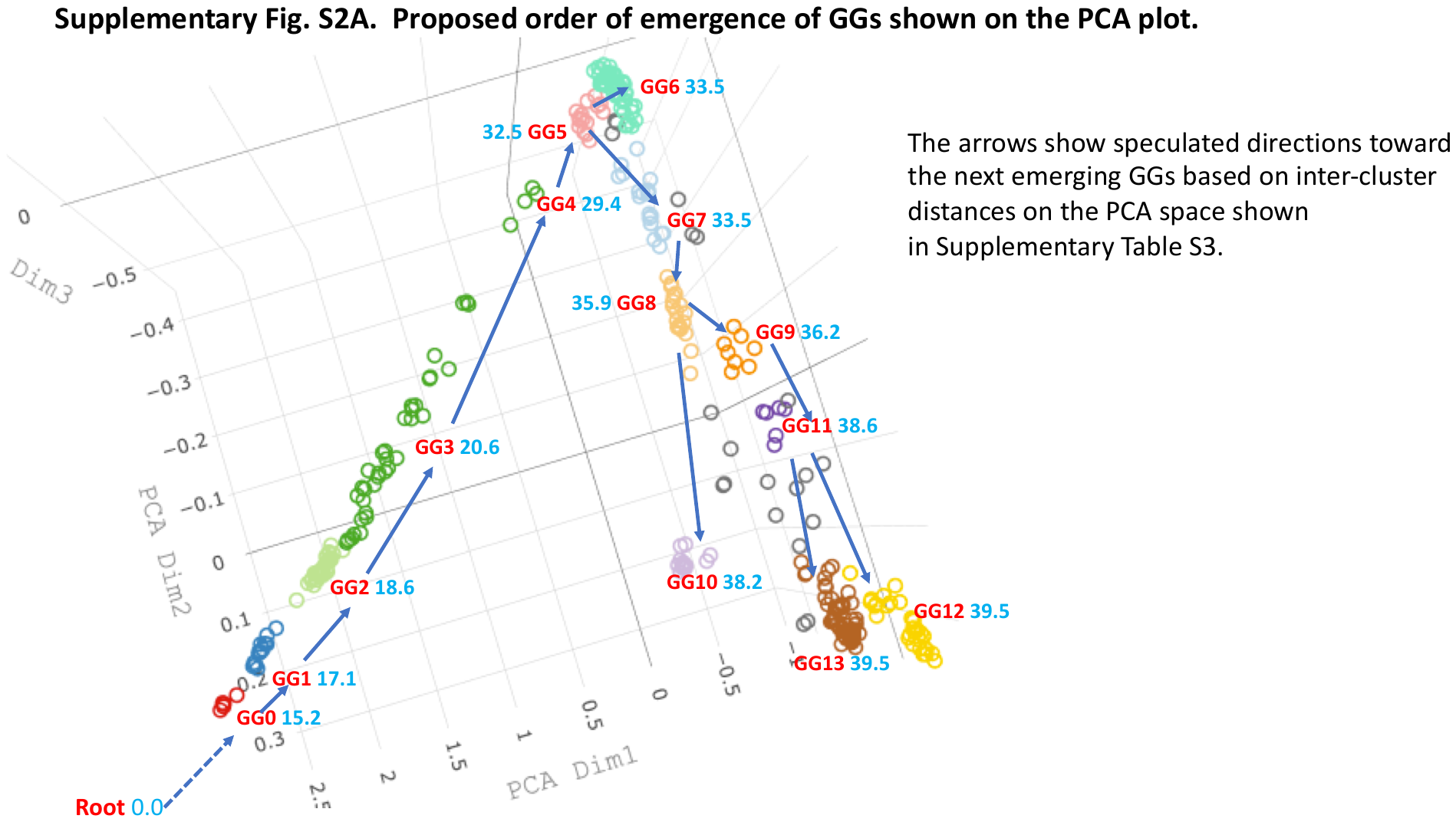

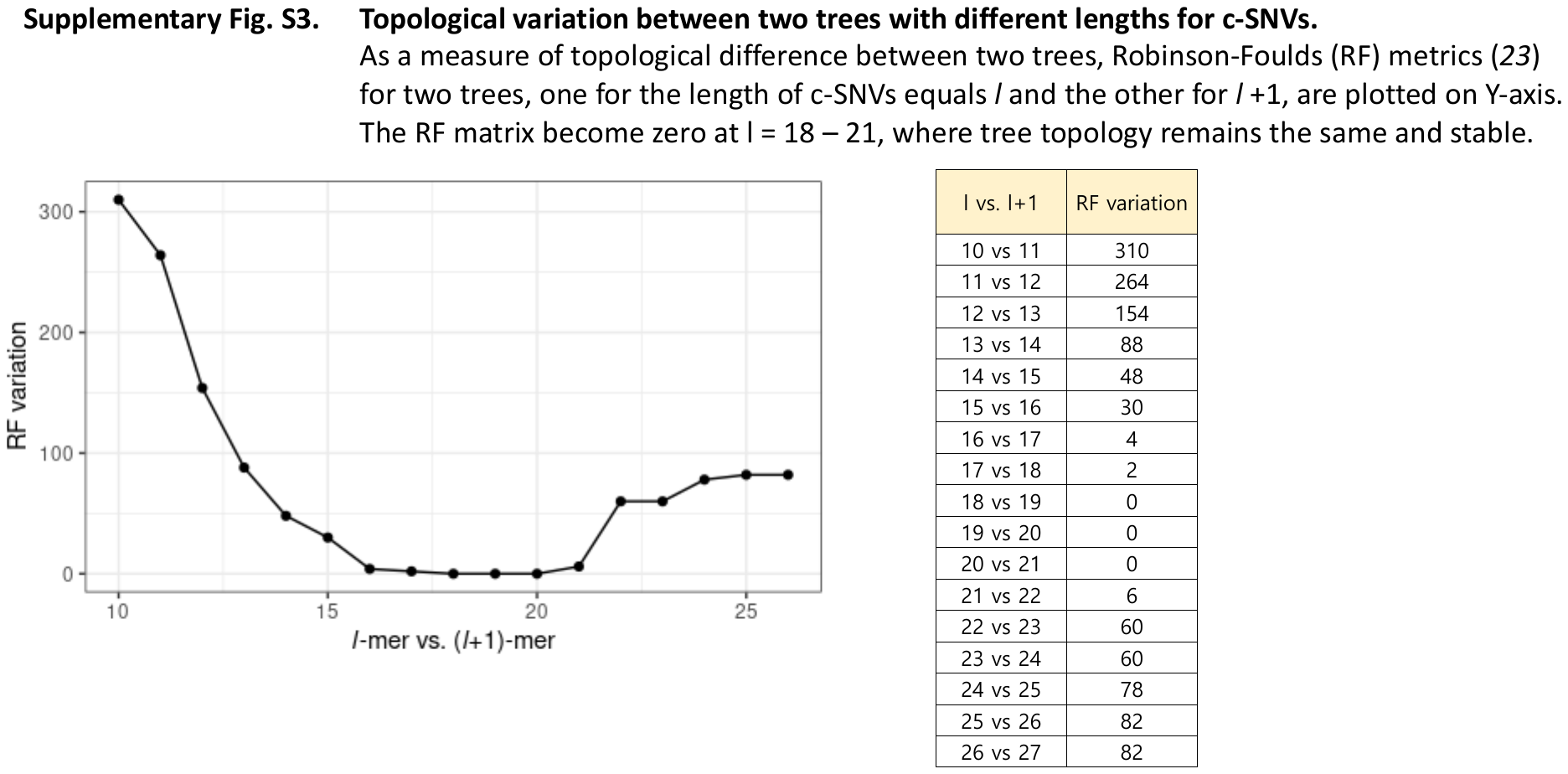

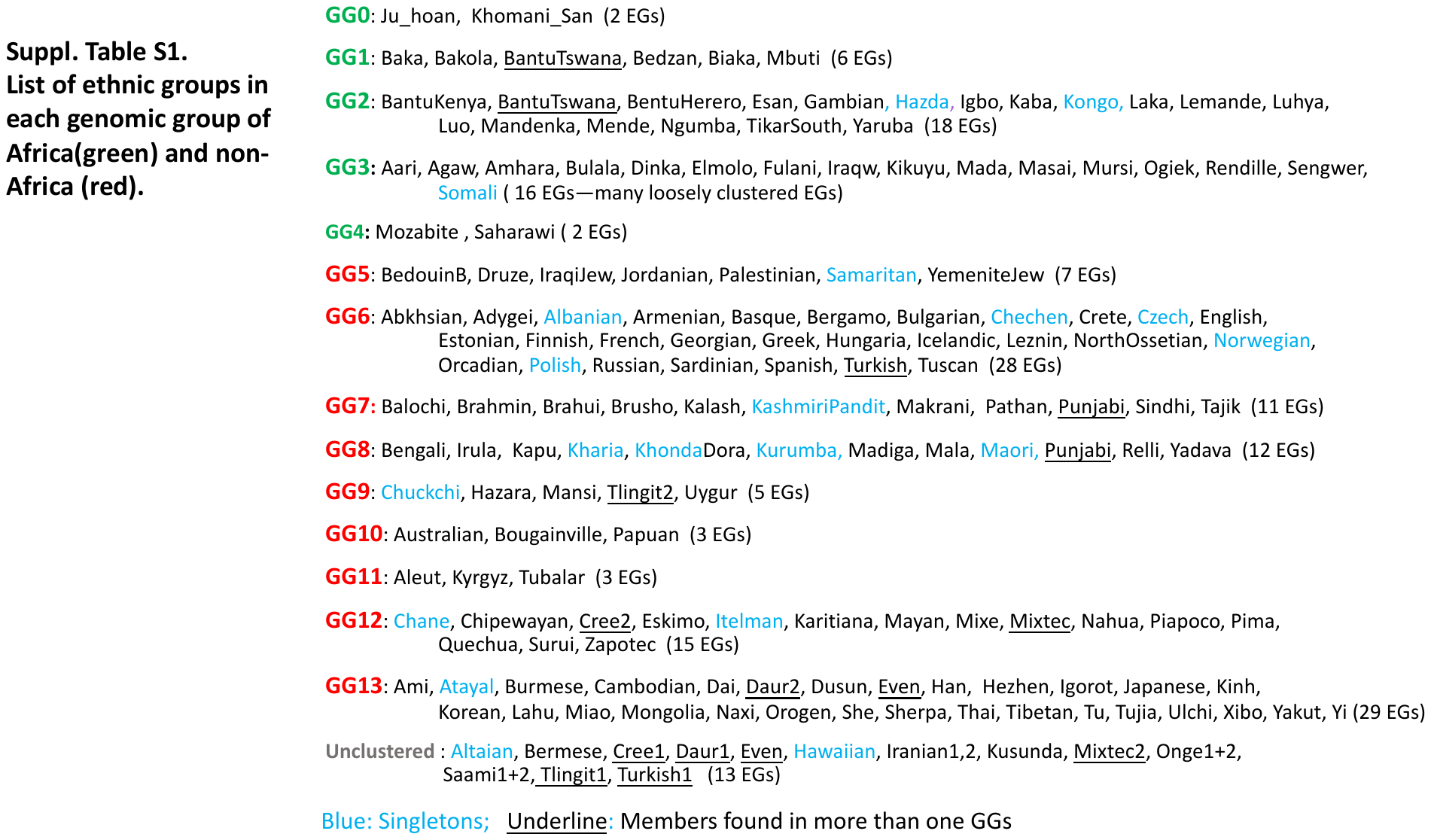

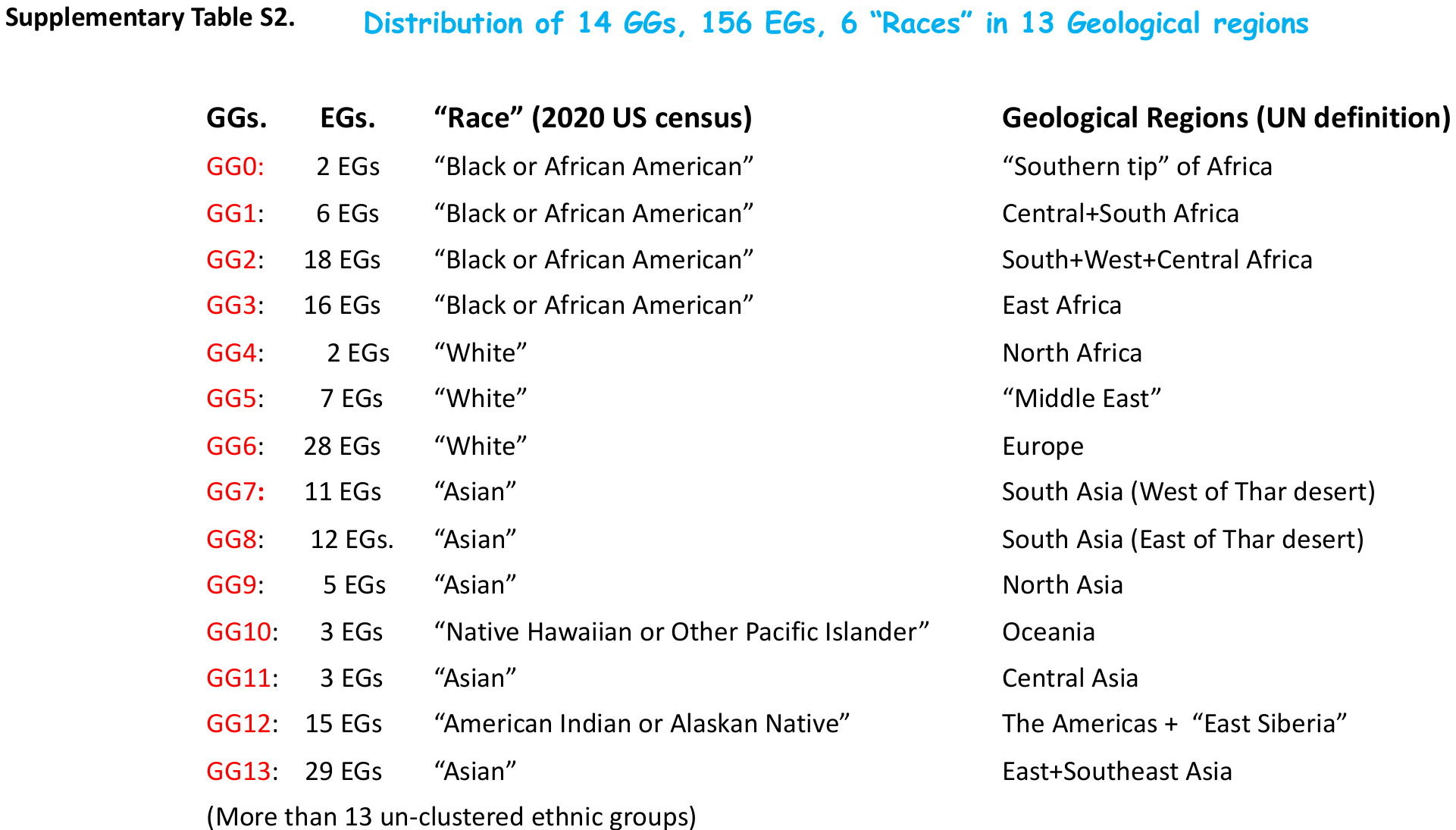

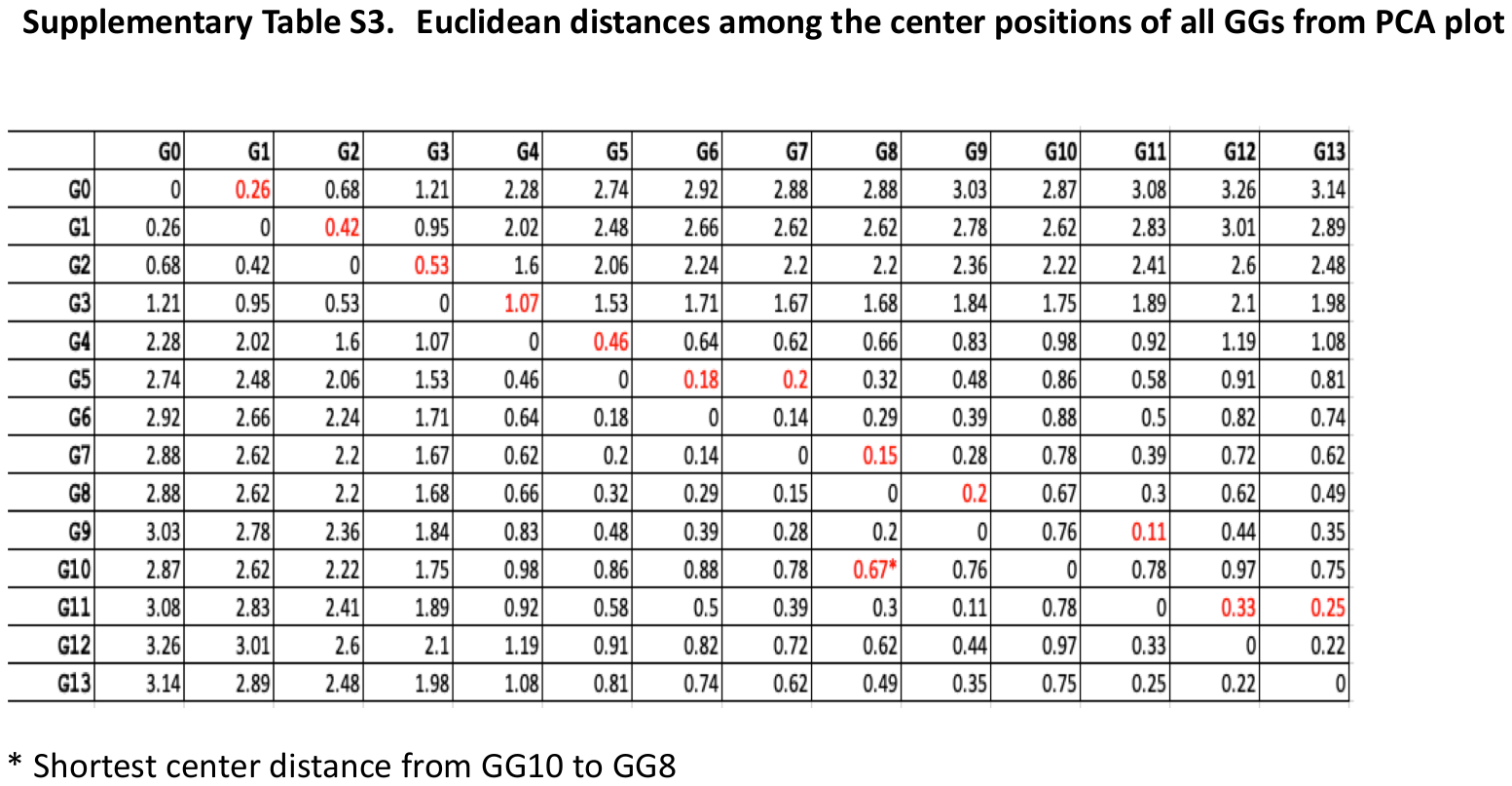
